# Fitness costs of herbicide resistance across natural populations of the common morning glory, *Ipomoea purpurea*

**DOI:** 10.1101/030833

**Authors:** Megan L. van Etten, Adam Kuester, Shu-Mei Chang, Regina S Baucom

**Affiliations:** Department of Ecology and Evolutionary Biology, University of Michigan, Ann Arbor, MI 48103; Plant Biology Department, University of Georgia, Athens, GA 30602

**Keywords:** cost of herbicide resistance, fitness cost, glyphosate, *Ipomoea purpurea*, germination, early life history, seed quality, trade-offs

## Abstract

Although fitness costs associated with plant defensive traits are widely expected, they are not universally detected, calling into question their generality. Here we examine the potential for life history trade-offs associated with herbicide resistance by examining seed germination, root growth, and above-ground growth across 43 naturally occurring populations of *Ipomoea purpurea* that vary in their resistance to RoundUp^®^, the most commonly used herbicide worldwide. We find evidence for life history trade-offs associated with all three traits; highly resistant populations had lower germination rates, shorter roots and smaller above-ground size. A visual exploration of the data indicated that the type of trade-off may differ among populations. Our results demonstrate that costs of adaptation may be present at stages other than simply the production of progeny in this agricultural weed. Additionally, the cumulative effect of costs at multiple life cycle stages can result in severe consequences to fitness when adapting to novel environments.

## Introduction

Plant defense is generally hypothesized to involve a cost. This expectation stems from the surprising observation of genetic variation underlying plant defense traits in many natural systems, whether the elicitor of damage is an herbivore, a pathogen, or an herbicide (Simms and Rausher 1987, 1989; Stahl et al. 1999; Baucom and Mauricio 2004; Bakker et al. 2006; Menchari et al. 2006; Délye et al. 2010; Kuester et al. 2015). If there were no costs associated with defense, traits conferring either resistance or tolerance to damage should increase to fixation rendering all individuals in the population highly defended (Rausher and Simms 1989). Despite our expectations of a trade-off between fitness and defense, however, reviews of the literature consistently show that costs are not ubiquitous regardless of the elicitor of selection or the study organism at hand (Bergelson and Purrington 1996; Coustau and Chevillon 2000).

Three main ideas have been proposed to explain the absence of such costs. First, there are a diverse number of potential mechanisms responsible for adaptation to a damaging agent, only some of which may incur a cost (Powles and Yu 2010; Vogwill et al. 2012). A single gene nucleotide substitution that leads to herbicide resistance, for example, may not alter the efficiency of translated proteins and therefore not incur a cost (e.g. Yu et al. 2007; Yu et al. 2010). On the other hand, a mechanism that provides resistance to a range of different herbicides through changes in growth may be more likely to impose fitness costs. Second, costs may not be detected if the genetic background is not properly controlled (Bergelson and Purrington 1996; Vila-Aiub et al. 2009b; Vila-Aiub et al. 2011). Control of the genetic background, either by performing crosses (Baucom and Mauricio 2004; Menchari et al. 2008; Giacomini et al. 2014) or ensuring replication across multiple genetic backgrounds (Cousens et al. 1997; Strauss et al. 2002) increases the likelihood that a cost will be detected (Bergelson and Purrington 1996). Third, researchers often examine only a portion of the life cycle (i.e., seed production or fecundity) and may do so in artificial and/or non-competitive conditions (Vila-Aiub et al. 2009b; Vila-Aiub et al. 2011). Studies that examine a range of traits are more likely to identify potential growth and/or fitness differences associated with plant defense compared to those that focus solely on measures of fecundity (Vila-Aiub et al. 2009b).

The phenomenon of herbicide resistance in plant weeds provides a particularly useful system to investigate the nature and types of costs associated with plant defense, since we know when selection by the herbicide began, the strength of selection, and often the frequency of herbicide use. However, as in other systems examining the evolution of plant defense, fitness costs of herbicide resistance are often not detected (Bergelson and Purrington 1996; Gemmill and Read 1998; Vila-Aiub et al. 2009b). Despite recommendations to control/increase the number of genetic backgrounds (Bergelson and Purrington 1996), and to examine multiple life history stages when determining if resistance incurs a cost (Primack and Kang 1989; Vila-Aiub et al. 2009b), only 25% of herbicide resistance studies control for background effects; further, only 7-10% of cost studies examine multiple stages of the life cycle (Vila-Aiub et al. 2009b). Fewer still examine the potential for fitness costs using a large number of naturally occurring populations sampled from a species’ range, an approach suggested almost 20 years ago (Cousens et al. 1997; Strauss et al. 2002). Just as the mechanism of resistance can vary among species, populations of the same weed have been shown to harbor different mechanisms of resistance to the same herbicide (Christopher et al. 1991; Christopher et al. 1992; Christopher et al. 1994; Preston and Powles 1998; Yu et al. 2008; Délye et al. 2010), thus increasing the likelihood that costs may likewise vary among populations. It is also possible, though rarely tested, that fitness costs have been ameliorated in some herbicide resistant populations relative to other populations due to the evolution of modifier loci (i.e. compensatory evolution, Darmency et al. 2015). The above hypotheses for the lack of costs are all interrelated: because resistance could be due to a variety of mechanisms (Délye et al. 2013a), costs may be apparent at only certain life history stages, expressed in particular environments (Vila-Aiub et al. 2009b), or apparent in some populations but not others. Thus, there remain crucial gaps in our understanding of where in the life history of a plant tradeoffs between fitness-enhancing traits and resistance might be apparent, and further, how ubiquitous such trade-offs may be across a species’ range (Vila-Aiub et al. 2011; Neve et al. 2014).

The common morning glory, *Ipomoea purpurea*, a noxious weed of US agriculture (Webster and MacDonald 2001), provides an excellent system to examine the strength and type of potential costs that may be present in natural populations. This species exhibits variability in resistance to glyphosate (Baucom and Mauricio 2008; Kuester et al. 2015), which is the main ingredient in the herbicide RoundUp^®^. RoundUp^®^ is currently the most widely used herbicide in agriculture (Fernandez-Cornejo et al. 2014), and of the approximately 30 resistant weeds that have been examined (Heap 2015), only a third are reported to express fitness costs (Ismail et al. 2002; Pedersen et al. 2007; Brabham et al. 2011; Giacomini et al. 2014; Shrestha et al. 2014; Vila-Aiub et al. 2014; Glettner and Stoltenberg 2015; Goh et al. 2015). *I. purpurea* has long been considered to exhibit low-level resistance to glyphosate (Culpepper (2006)), and previously we have shown that this low-level resistance (estimated as proportion leaf damage) has an additive genetic basis and is under positive selection in the presence of the herbicide (Baucom and Mauricio 2008). Further, a recent replicated dose-response experiment of 43 populations sampled from the southeastern and Midwest US showed that some populations of *I. purpurea* exhibit ~100% survival after application of the field dose of RoundUp^®^ (*i.e*., resistance), whereas other populations exhibit high susceptibility (Kuester et al. 2015). Although we find variability in resistance across natural populations, it is unclear if this defense trait involves a cost. We investigated this question within one population using artificial selection for increased/decreased resistance and discovered that the seed production of individuals from the increased resistance lines was not significantly lower than that of susceptible lines in the absence of the herbicide, suggesting that there may not be a fecundity cost associated with resistance in this species. However, there was some indication that progeny quality may be lower in resistant individuals - resistant lines exhibited a trend for reduced seed viability compared to susceptible lines (Debban et al. 2015). This finding suggests that trade-offs between fitness enhancing traits (e.g., germination and resistance) may be present within this species, which could manifest as a cost by reducing the overall fitness of resistant compared to susceptible lineages in the absence of herbicide.

Here we determine if there are trade-offs associated with resistance by examining germination, early root growth and above-ground growth across 43 populations of *I. purpurea*. We specifically ask the following: (1) are there potential trade-offs associated with resistance across this species’ range in the US, manifest in the form of (i) lower germination and/or (ii) smaller size at early life history stages (i.e., early germinant, young plant)?, and (2) do resistant populations exhibit the same type of potential tradeoff, which may indicate the nature and expression of fitness costs may vary across populations?

## Materials and Methods

### Seed collection and control of maternal/environmental effects

Multiple fruits were collected from up to 79 individuals separated by at least 2 m from 43 populations located across the Midwest and Southeastern US (Table S1; Fig S1). These seeds (hereafter field-collected seeds) were used in several experiments to determine resistance, germination and early growth characteristics. To homogenize the effects of maternal environment on seed quality, we chose a subset of the populations (N=18), grew them in a common greenhouse for one generation and collected the autonomously self-pollinated seeds from a similar growing and mating system environment (hereafter once-selfed seeds).

### Estimate of herbicide resistance

To determine glyphosate resistance across populations, a dose-response experiment was conducted by planting a single field-collected seed from 10 randomly chosen maternal lines from each population in six glyphosate treatments (including a non-herbicide control treatment) in each of two greenhouse rooms. Full details of the dose-response experiment are presented in Kuester *et al*. (2015) - for simplicity, we present resistance as the percent survival per population at 1.70 kg a.i./ha of glyphosate, a rate which is slightly higher than the suggested field rate of 1.54 kg a.i./ha. Individual seeds were scarified, planted, allowed to grow for three weeks, and then treated with the herbicide (PowerMax Roundup; Monsanto, St. Louis, Missouri) using a hand-held CO_2_ pressurized sprayer (Spraying Systems Co., Wheaton, IL). Survival was scored three weeks after treatment application, and the population estimate of resistance was determined as the proportion of individuals that survived glyphosate.

### Germination

We performed three germination experiments to determine if resistance influenced seed traits. First, we examined germination using field-collected seeds in a petri-dish assay in the laboratory; second, we examined germination of the field-collected seeds in the soil in the greenhouse; and third we performed a petri-dish assay in the lab using seeds generated via selfing in the greenhouse (once-selfed seeds) to examine the potential for maternal field environmental effects. For the first experiment using field-collected seeds, we measured seed weight and germination characteristics using field-collected seeds from each population (N=43). Up to five (ave 4.6) seeds from 8-79 maternal lines per population (ave 38, total 1621, see Table S1 for exact sample sizes per population) were randomly chosen for the germination test. From this pool of seeds we randomly chose a subset of families per population (8-49 maternal lines per population; Table S1) for which the selected seeds were weighed (as a group) to determine the average seed weight. All of the selected seeds were placed in a small petri dish (one dish per family), submerged in filtered water and allowed to germinate in the lab under ambient light and temperature. Water was added as necessary every three days to prevent drying out. Petri dishes were completely randomized across lab benches. Germination was scored periodically until no further germination was recorded. Final pre-scarification germination was scored after 16 days, with successful germination considered the emergence of a normal radicle. At this time, seeds that had not imbibed water (by visual determination) were scarified and germination was again scored after 1 week. We recorded the final number of seeds exhibiting normal germination, the number of seeds needing scarification, the number of scarified seeds that germinated, and the number that had abnormal germination. For the second germination assay, we examined germination data from one replicate (housed in a single greenhouse room) of the dose-response experiment mentioned earlier in which seeds were scarified and planted in conetainers with 1 seed per pot (10 maternal families per population). Germination was scored after three weeks and used to calculate the percentage of seeds that germinated.

For the third and final germination assay, we used seeds from maternal lines that were selfed once in the greenhouse. Two sets of five seeds for up to 8 maternal lines (randomly selected) for each of 18 populations were placed in petri dishes with water. Germination was scored after 11 days. If seeds had not imbibed water they were scarified and scored again in one week. We calculated the percentage of seeds with normal germination prior to scarification, the percentage of seeds that germinated after scarification, and the percentage that had abnormal germination.

### Early root and above-ground growth

To examine early root growth, we again used the once-selfed seeds and measured root length four days after the germination assay began. We chose to first scarify the seeds in this assay to standardize water absorption among individuals. Two sets of five seeds for up to 8 maternal lines (randomly selected) for each of 18 populations were scarified and placed in petri dishes with water. Germination was scored after 1, 4 and 7 days. On day 4 petri dishes were scanned and the root length was measured using Image J (Abramoff et al. 2004) for each germinated seed.

We next examined early growth traits of greenhouse-grown individuals to determine if there was a relationship between resistance and plant size (i.e., are plants from resistant populations smaller?). To do so we used measurements from plants from the dose-response experiment prior to herbicide application. Three weeks after planting (and prior to spraying) we measured the height of the stem, the number of leaves and length of the largest leaf on each individual planted per treatment per population (total N=2908, Table S1 for exact sample sizes per population).

### Statistical analysis

*Field-collected seeds*—We assessed the relationship between resistance and progeny quality using mixed model analyses of variance. We used a generalized linear mixed-effect model to examine final germination, germination before scarification, abnormal germination, seeds needing scarification, and germination after scarification with resistance and population (random) as predictors using the glmer function in the R package lme4 with a binomial distribution. All of the binary measures were coded as 1 or 0. Seed weight (g) was modeled using a mixed model with resistance and population (random) as predictors using the lmer function in the R package lme4. Additionally, previous studies have indicated a geographic pattern of resistance in this species (Kuester et al. 2015). To ensure that the above results were not an artifact of geography, we added latitude and longitude (scaled) of the population in the above models. For the experiment examining germination in soil, we modeled germination with resistance level and population (random) as predictors using a binomial distribution.

*Once-selfed* seeds—Similar to the field-collected seeds, we used mixed-model binomial regressions to assess the effect of resistance on germination characteristics of the once-selfed, greenhouse generated seeds. We modeled germination before scarification and germination after scarification with resistance and population (random) as predictors using a binomial model. To determine if the maternal environment in which the seeds developed influenced germination, in a separate model we compared germination between maternal environments (i.e., field collected seeds versus seeds propagated in the greenhouse) by including maternal environment as a treatment effect in the model. To do so, we modeled final germination using treatment, resistance, population (random) and treatment*resistance as predictors using a binomial distribution. An interaction between treatment and resistance would indicate that the maternal environment differently influenced germination.

*Early growth and size*—We next used mixed model analyses of variance to determine if more resistant populations exhibited early growth life-history trade-offs. We separately considered root length of the early germinant and plant size. We examined root length using the once-selfed seeds in two different models. The first and more basic model examined the influence of resistance and population (random) on log-transformed root length (cm) 4 days post germination. A difference in root length, however, could be due to differences in either growth rate of the radicle or differences due to the timing of germination, *i.e*., when growth began following germination. To distinguish between these two potential explanations, we calculated an estimate of germination speed, the time to 50% germination - a shorter time would suggest that seeds began growing sooner after water was added. We used the germination data from days 1, 4, and 7 to obtain a population level estimate of the time to 50% germination using a germination Hill function (El-Kassaby et al. 2008). This function decomposes germination into 4 parameters: a, the germination capacity; b, the steepness of the curve; c, the time to 50% germination; and y_0_, the lag time before germination. We used the nonlinear least squares (nls) function in R to estimate the b and c parameters. We chose to pool the data on a population level to increase the accuracy of the estimation. The time to 50% germination (c) was then used as a covariate in the more complex model of root length that included resistance, population (random) and time to 50% germination.

We next examined height (cm), leaf number and leaf size (cm) of plants grown from field-collected seeds (~3 weeks growth in greenhouse) to determine if resistance incurs early growth life-history trade-offs. We used each trait in separate mixed models with replicate, rack within replicate (random), resistance and population (random) as predictors. Residuals of leaf size were not normal so a box-cox transformation (λ = 2.0) was used to achieve better fit.

Finally, we performed a Principle Components Analysis (PCA) using the population averages of several traits from the field-collected seeds, which included seed weight, germination percentage, percentage of abnormally germinating seeds, percentage of successfully germinating scarified seeds, early plant height, leaf number and leaf size to visually examine the data and determine how populations differed along the two axes retained. This analysis was performed using PROC FACTOR in SAS with a varimax rotation to obtain more easily interpretable axes. Loadings and the proportion variance explained for each factor with an eigenvalue >1 can be found in Table S3.

## Results

### Germination

We found a strong and significant negative relationship between resistance and the percentage of field-collected seeds that germinated (Fig. 1a). This is true both of seeds that germinated before scarification (β = -4.93, χ^21^ = 24.66 P < 0.0001) and the total number that germinated (including those that germinated after being scarified; β = -5.20, χ^21^ = 24.80, P < 0.0001; Fig. 1a). In addition to a decline in germination, several other measures of seed quality also declined with increasing resistance. We found a higher percentage of abnormally germinating seeds (β = 4.24, χ^21^ = 33.20, P < 0.0001) in that, instead of exhibiting normal germination, a non-viable embryo would be ejected from the seed coat with no further growth. Furthermore, some seeds simply did not imbibe water; we scarified these seeds to determine if they were viable but potentially dormant. Populations with greater resistance had more seeds that needed scarification (β = 1.52, χ^21^ = 5.00, P = 0.03), of which fewer seeds that subsequently germinated (β = -5.50, χ^21^ = 8.81, P = 0.003). We also found that populations with higher resistance produced lighter seeds (β = -0.005, χ^21^ = 4.69, P = 0.03), indicating that resistance influenced multiple measures of seed quality for seeds collected from the field. All of these relationships remain significant after accounting for longitude and latitude of the populations except for the percentage needing scarification (Table S2), suggesting that the patterns we find are not due to a simple geographic pattern. We similarly uncovered a negative relationship between germination and resistance when seeds from these populations were planted in soil in the greenhouse (β = -0.79, χ^21^ = 16.09, P < 0.0001; Fig. 1b).

**Fig 1.**
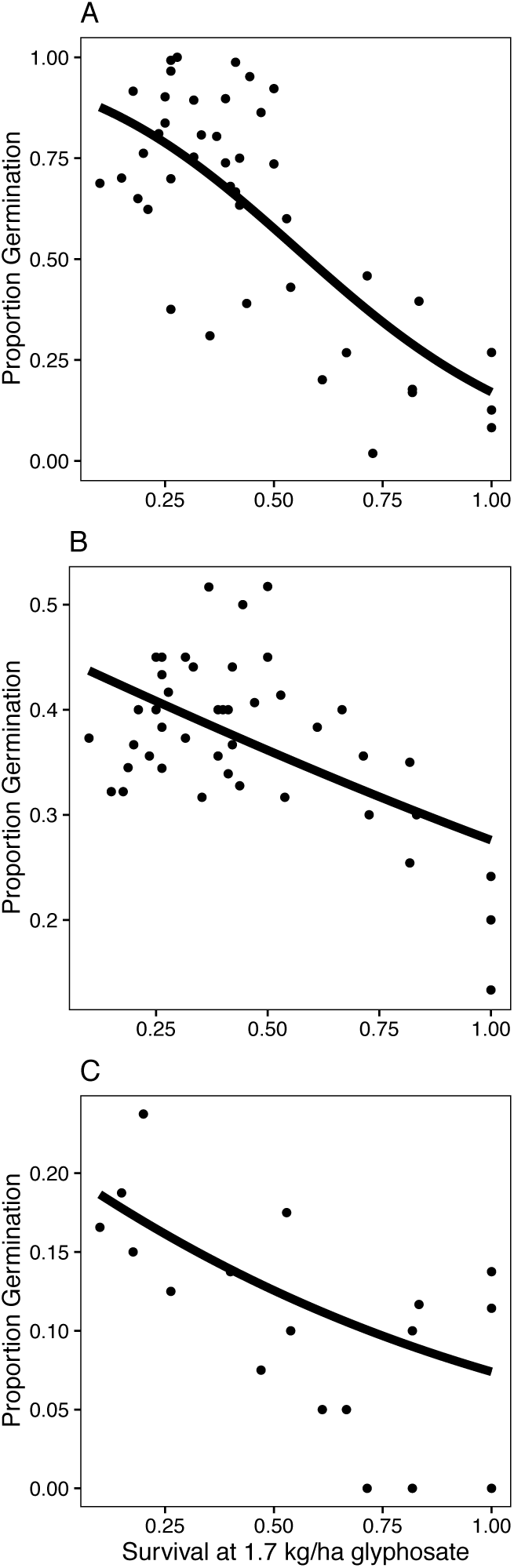
Relationship between herbicide resistance and proportion germination for (a) field-collected seeds in petri dishes, (b) field-collected seeds in soil, and (c) once-selfed seeds (note the differences in the y-axis scale). Points are the mean per population, lines are the average marginal predicted probabilities from the appropriate model.

The negative relationship between resistance and germination was supported by the results from the once-selfed seeds grown in a common environment for a generation (Fig. 1c). Prior to scarification, very few of the greenhouse-grown seeds imbibed water and germinated (2.0%) and there was no effect of resistance (β = 184.2, χ^21^ = 0.81, P = 0.37). After scarification, however, there was a significant negative relationship between germination and resistance (β = -1.18, χ^21^ = 6.42, P = 0.01, Fig. 1c). This effect remained significant after accounting for latitude and longitude (β = -1.13, χ^21^ = 5.49, P = 0.02). Interestingly, the decrease in germination of once-selfed, greenhouse-generated seeds was significantly less than the field collected seeds (treatment * resistance: P = 2.14, χ^21^ = 27.8, P < 0.0001) suggesting that maternal environmental conditions influence the quality of seeds produced. In addition, we found a much lower rate of abnormal germination in the once-selfed seeds (~10%) compared to the field collected seeds, and the level of abnormal germination showed no relationship with resistance (β = 0.31, χ^21^ = 0.23, P = 0.63). These results suggest that, while germination costs are consistently detected between experiments in which the maternal environment differed, field environmental conditions exacerbate the strength of the germination cost.

### Early root and aboveground growth

To test whether growth differed according to resistance status, we scarified and germinated the once-selfed seeds then measured root growth after 4 days. There was a much higher germination rate of these seeds (86%) compared to the previous experiment (2% pre-scarification) and the majority occurred before day 4. While there was a nearly significant negative relationship between root length and resistance (β = -0.38, χ^21^ = 3.19, P = 0.07; Fig. 2a), including the time to 50% germination in the model removed this effect (resistance: β = -0.07, χ^21^ = 0.09, P = 0.76; time to 50% germination: β = -0.37, χ^21^ = 4.85, P = 0.03), suggesting that the difference in root length was due to the timing of germination rather than a difference in growth rate. A difference in plant size was also found in the 3-4 week old plants grown in soil from the field-collected seeds - plants from more resistant populations had smaller above-ground structures on average than plants from less resistant populations (height: β = -7.42, χ^21^ = 7.20, P = 0.007; leaf number: β = -1.11, χ^21^ = 15.86, P < 0.0001; largest leaf width: β = -4.97, χ^21^ = 17.32, P < 0.0001; Fig. 2b-d).

**Fig 2.**
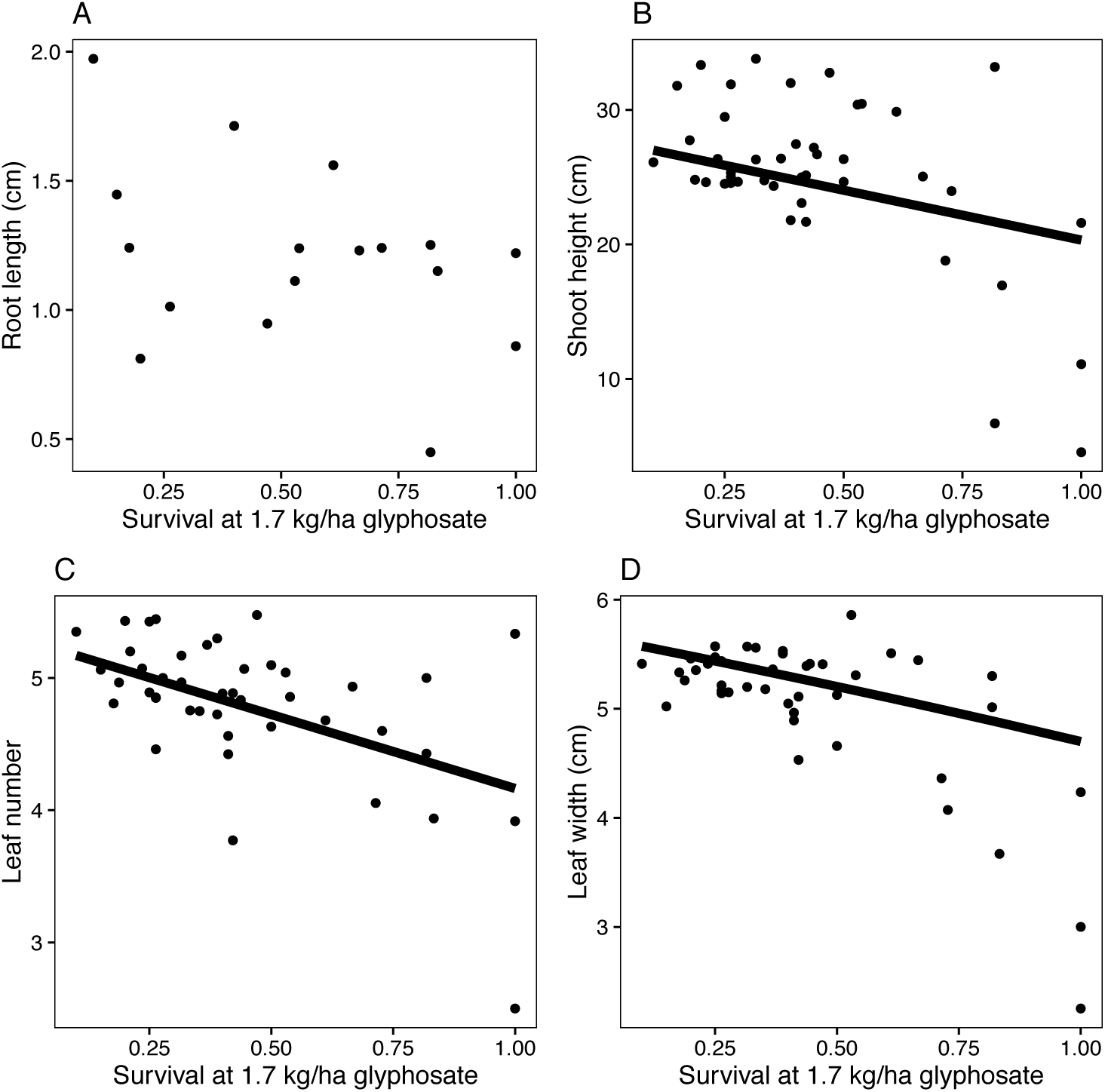
Relationship between herbicide resistance and (a) root growth after 4 days, (b) shoot height (c) leaf number and (d) width of the largest leaf after 3 weeks. Points are the mean per population and lines are the average marginal predicted probabilities from the appropriate model.

### Visualization of cost-related traits

We next examined germination and early growth traits from the original field-collected seeds using a principle components analysis (PCA) to determine if there was variation among populations in the expression of cost-related traits (full results Table S3). The first 3 principle components of the PCA explained 77% of the variance, with the first principle component (PC) loading with the early growth traits while the second loaded with seed traits and the third with the proportion of seeds that required scarification to successfully germinate. Populations with higher resistance scored lower on PC1 (b = -2.22, r^2^ = 0.28, t_41_ = -4.03, P = 0.0002) and PC2 (b = -2.12, r^2^ = 0.26, t_41_ = -3.79, P = 0.0005), but not on PC3 (b = -1.02, r^2^ = 0.06, t_41_ = -1.61, P = 0.12). Using the first two PCs to plot the results, resistant populations occur mostly in the lower left quadrant (smaller plants, lighter seeds, lower germination and more abnormally germinating seeds) and have a wider spread than less resistant populations (Fig. 3). Furthermore, while resistant populations scored lower on PC1 and PC2 in comparison to susceptible populations, some resistant populations exhibited early growth traits that were similar to susceptible populations and yet scored very low on germination traits (e.g., pop num 5) whereas other resistant populations exhibited similar germination traits compared to the susceptible populations, but were smaller in stature than susceptible populations (e.g., pop num 51). Thus, it appears that the type of cost may vary among populations sampled from North America.

**Fig 3.**
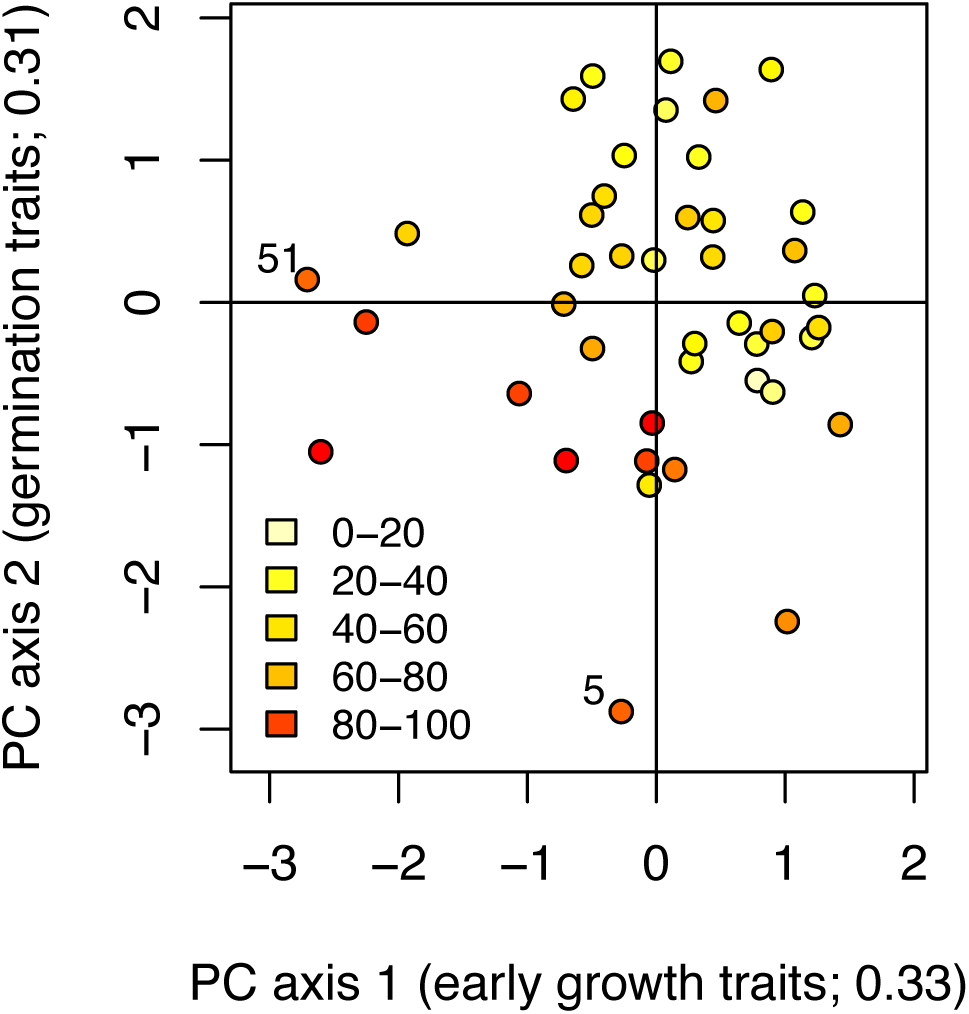
Scatter plot of PCA results showing average principle components axis 1 (higher values indicate larger plants) and principle components axis 2 (higher values indicate heavier seeds, higher germination, and fewer abnormally germinating seeds) values for field-collected seeds with circle color indicating survival at 1.7 kg/ha glyphosate. The proportion variation explained by each axis is noted in axis labels.

## Discussion

Here we show that glyphosate resistant populations of the common morning glory exhibit life-history trade-offs associated with resistance, and, that these trade-offs may vary among populations. Our series of experiments uncovered three notable findings: First, we found a negative linear relationship between germination and resistance indicating that resistant populations have a lower germination rate than susceptible populations. This negative relationship persisted when using seeds generated from a common greenhouse environment showing that this result is not due solely to field environmental and/or maternal effects. Second, we found that individuals from resistant populations were smaller than individuals from susceptible populations, indicating that resistance influences early plant growth. Interestingly, we also found evidence that the two types of trade-off may differ among populations—using PCA, we show that some resistant populations produce normally sized plants, but score low on germination traits, and *vice versa*. Below, we detail how these results add further strength to the suggestion that a variety of life stages and populations sampled across the species’ range should be assessed when testing the hypothesis that resistance incurs a fitness cost (Délye et al. 2013a).

### Fitness costs: seed germination and early plant size

It is difficult to determine how common germination differences associated with resistance may be among weeds since many studies focus on seed quantity rather than seed quality. There is some indication that germination may be affected in other glyphosate resistant species. Dinelli et al. (2013) found reduced germination of glyphosate resistant *Ambrosia trifida* populations, while Ismail et al. (2002) found greater germination of resistant biotypes of goosegrass (*Eleusine indica*). More broadly, life history trade-offs may be specific to the herbicide and/or species in question or the type of mutation conferring resistance (O'Donovan et al. 1999; Vila-Aiub et al. 2005; Délye et al. 2013b). For example, only one of two different resistance mutations in ACCase resistant *Lolium rigidum* had more stringent germination requirements (seeds germinated poorly in the dark and required fluctuating temperatures to break dormancy) than the susceptible genotype (Vila-Aiub et al. 2005). Similarly, Délye *et al*. (2013b) found differential effects on germination among resistance mutations to ACCase in *Alopecurus myosuroides*. Both of these studies report that the resistance mutation led to delayed germination. Such a delay in germination may affect fitness, especially in agricultural settings where germinating too early can lead to removal by pre-sowing practices and germinating too late can lead to intensified competition with already established plants (Weaver and Cavers 1979; Barrett 1983; Mortimer 1997; Forcella et al. 2000; Owen et al. 2014). Our analysis of root growth suggests that differences in plant size in *I. purpurea* may be due to a similar delay in germination in resistant populations.

The reduced growth of resistant compared to susceptible populations that we uncovered could lead to decreased competitive ability and subsequent lower fitness in the presence of competition if, as has been found in other herbicide resistant weeds, the difference in growth persists to adult plants (Weaver and Warwick 1982; Ahrens and Stoller 1983; Holt 1988; Alcocer-Ruthling et al. 1992; Williams et al. 1995; Vila-Aiub et al. 2005; Tardif et al. 2006; Vila-Aiub et al. 2009a). This type of life-history trade-off, which ultimately may manifest as a fitness cost, is also likely to be species, mutation and environment specific. For example, *Lolium rigidum* has evolved herbicide resistance via a variety of mutations ranging from target site (Christopher et al. 1992; Yu et al. 2008) to non-target site (Christopher et al. 1991; Christopher et al. 1994; Preston and Powles 1998). Target site mutations in the acetohydroxyacid synthase gene result in little cost in growth (Yu et al. 2010). On the other hand, herbicide resistance mediated by the cytochrome P450 complex resulted in reduced biomass and decreased competitive ability (Vila-Aiub et al. 2009a).

An alternative explanation for the decline in germination and growth we identify using the field-collected seeds is that some other co-varying population characteristic such as soil fertility, spraying regime, herbivore levels, or the many other biotic and abiotic factors that can influence seed development differed among resistant and susceptible populations (Roach and Wulff 1987; Fenner 1991; Schmitt et al. 1992; Platenkamp and Shaw 1993; Galloway 2001). These differences may explain the stronger decline in germination in the field-collected seeds compared to the once-selfed seeds. Several lines of evidence, however, suggest the relationship between resistance and germination that we uncovered represents a true fitness cost rather than simply an effect of the environment (e.g., driven by maternal effects, latitude, spray environment). First, the relationship between resistance and germination appears approximately linear, which is expected as the frequency of resistant individuals increases. If the trade-offs identified herein were due simply to glyphosate exposure, with resistant populations exhibiting abnormal seed development after surviving glyphosate application, we would expect to see a binary distribution of seed quality of populations that had been sprayed and those that had not, rather than a linear trend with resistance. Second, the negative relationship between resistance and germination is maintained after a generation in a common greenhouse environment—an effect should disappear if the decrease in fitness was due to glyphosate exposure in the field. Finally, our results parallel those from a recent experiment that specifically controlled for genetic background and environmental effects using *I. purpurea* plants from a single population (Debban et al. 2015). Individuals from this population were artificially selected for increased or decreased resistance for three generations under controlled greenhouse conditions, and, similar to results presented here, the increased resistance lines had a larger percentage of “bad” seeds that ejected the embryo. That the results from one population utilizing a controlled genetic background are mirrored across many populations collected from the landscape provides strong evidence that lower germination represents a fitness cost of glyphosate resistance in this weed species.

While we detected lower germination in herbicide resistant populations across multiple experiments suggesting a true trade-off, we also found differences between experiments in the strength of the relationship. This suggests both an underlying genetic basis as well as an environmental component influence the expression of the trade-off. Compared to the field-collected, unscarified seeds, the once-selfed, unscarified seeds (Fig. 1) had a low germination rate suggesting that the seed coat was perhaps more pristine in seeds generated in the greenhouse. However, if scarified, we found that the once-selfed seeds exhibited high germination and the relationship between resistance and germination remained negative indicating a fitness decline associated with resistance.

The physical seed coat is the primary mechanism of dormancy in this species (Brechu-Franco et al. 2000); thus, environmentally-induced physical differences in the seed coat (e.g. thickness or waxiness) or its degree of degradation (e.g. mechanical disruption or seed storage differences) likely influences germination timing. Although it is clear that the environment influences seed germination in this species, that we consistently observed a decline in germination with resistance across multiple experiments suggests that there is an underlying genetic basis to the cost of resistance.

Another striking difference between experiments was in the frequency of abnormal seeds produced. The once-selfed seeds had almost no abnormal germination (i.e., no dead embryos that were ejected from the seed coat) while some of the field-collected populations had a high level of abnormal germination. In fact, the strong decline in germination for field-collected seeds was due primarily to this abnormal germination. Abnormal germination could be due to a variety of environmental causes (e.g. herbicide application, nutrient availability, competition, etc) or be a cost of resistance that is only induced under field conditions. Our results suggest that in a benign environment, such as the greenhouse, the seeds in general are of high quality (high germination, fewer abnormal germinants) but there is a cost of resistance that increases the time it takes to germinate (based on root growth experiment), possibly leading to smaller plants at any given point. On the other hand, under field conditions, populations with higher resistance produce more abnormal seeds (due to either environmental differences or an environmentally induced cost of resistance) and the normal seeds may still have an increase in the time it takes to germinate leading to smaller plants at any given point.

Interestingly, by visually examining the germination and growth traits in a PCA, we find variation in the type of potential cost among resistant populations. While some resistant populations fell into the “poor germination” axis, other resistant populations fell into the “poor growth” axis compared to susceptible populations. There are at least three possible reasons for this difference: different resistance genes, different compensatory mutations or different genetic backgrounds. First, the gene(s) involved in resistance may vary among populations leading to different costs. Independent origins of resistance to herbicide have been found in other species (Délye et al. 2010) and these different mutations often incur different fitness costs (Vila-Aiub et al. 2005; Délye et al. 2013b). Second, the resistance gene(s) may be the same amongst populations but each population may have different compensatory mutations that lead to different costs (Darmency et al. 2015). Third, the resistance gene(s) may behave differently in different genetic backgrounds (Paris et al. 2008). These distinctions are important because they would differentially affect the evolutionary trajectory of herbicide resistance. For example, if populations differ in the gene(s) involved, each population may have a very different set of costs, benefits and evolutionary trajectories, which would need to be incorporated in models.

It is currently unknown if the trait trade-offs identified here are pleiotropic or due to linkage to the resistance gene. The most restrictive definition of a cost requires that the decrease in fitness is due to the resistance allele itself - either the actual allele or through it acting pleiotropically (Bergelson and Purrington 1996). Given that we do not know the identity of the loci involved in either resistance or the abnormal germination and reduced growth, we cannot entirely rule out physical linkage between resistance genes and cost genes, in which case the “cost” could quickly become unlinked over generations (Lewontin 1974; Hartl and Clark 1989). For some species, easily identifiable mutations in the enzyme targeted by the herbicide can be linked to resistance, i.e. target site resistance (TSR). However, preliminary work suggests that glyphosate resistance in *I. purpurea* is due to non-target site mechanism (NTSR; Leslie and Baucom, *unpublished data*), and as such elucidating the genetic basis of both resistance and the cost will be a non-trivial endeavor. Furthermore, it is rare that genes underlying costs are identified; most documented cases of the genes involved in the cost of resistance is when TSR mutations lead to poor performance of the enzyme on its natural substrate (Vila-Aiub et al. 2009b). As far as we are aware, no study has identified the genes involved in the cost of resistance when the mechanism of resistance is NTSR. One intriguing possibility for this species stems from a previous study that compared transcript expression levels of artificially selected lines of resistant and susceptible *I. purpurea* plants following herbicide application (Leslie and Baucom 2014). One of the differences between the replicated resistant and susceptible lines was a lower expression of pectin methylesterase (PME) in the resistant plants. This enzyme has been shown to play a role in breaking seed dormancy (Ren and Kermode 2000) and stem elongation (Pilling et al. 2000). Thus, the decreased expression of PME in resistant plants may explain both the reduced germination and growth in populations with higher resistance.

### How might life history trade-offs influence the evolutionary trajectory of resistance in this species?

Recent work in this system has shown that populations of the common morning glory sampled from 2012 exhibit higher levels of resistance compared to the same populations sampled in 2003 (Kuester et al, *In* Review). Interestingly, however, the difference in resistance between sampling years was only slight, i.e., 62% survival at 1.7 kg ai/ha in 2012 vs 57% survival in 2003 samples. It is possible that the life-history differences that we identified here are responsible, at least in part, for maintaining resistance between sampling years. For example, the lower germination of resistant types would manifest as a fitness cost if resistant and susceptible types produce approximately the same number of total seeds (or if R < S); if, however, resistant types produce enough viable seed to offset the lowered germination, then overall fitness would not be impacted and resistant types would not be at a relative disadvantage. While we have not examined seed production across all 43 populations examined herein, a common garden study of glyphosate susceptible and resistant families of this species found there was no difference in total seed production of resistant compared to susceptible lines (Debban et al 2015), indicating there is no cost of resistance in terms of seed quantity. That we similarly find poor germination between these experiments and those using genetic lines developed from a single population strongly supports the finding that germination quality is a true fitness cost of glyphosate resistance in this species. Further, the differences in growth that we have detected between resistant and susceptible populations could potentially manifest as a fitness cost if in competition, an effect which remains to be tested in this system.

In summary, we found reductions in seed quality across replicated herbicide resistant populations of the common morning glory. Although most studies use seed quantity as a proxy for fitness, our results highlight that reductions in progeny quality are an equally, if not more, important cost of adaptation in *I. purpurea*. Given that fitness costs are thought to arise from a variety of mechanisms (allocation of resources, ecological costs, etc.), our results suggest that a high priority should be placed on the examination of multiple stages of the life cycle when assessing potential costs and not just seed quantity. Furthermore, because the strength of this cost could maintain the efficacy of a globally important herbicide, this work illustrates the utility and importance of integrating evolutionary principles into management scenarios (Gould 1995).

## Acknowledgements

We thank E. Fall, A Wilson, A Jankowiak, D York, N Gabry, S Sanchez for assistance and J Vandermeer, MA Duffy, and PJ Tranel for comments on earlier drafts of the manuscript. This work was funded by USDA NIFA grants 04180 and 07191 to RSB and SMC.

## Supporting information

**Table S1.** Population characteristics including population number, state, crop type, latitude and longitude and sample sizes for experiments with field-collected seeds (N_fam_ germ = number of families in the germination assay for field collected seeds; N_seeds_ germ = number of seeds in the germination assay for field collected seeds; N early growth = number of seeds in the early growth experiment, N_fam_ once selfed = number of families in the germination assays for the once-selfed seeds).

| Population number | State | Crop | Latitude | Longitude | $N_{\text{fam germ}}$ | $N_{\text{seeds germ}}$ | $N_{\text{early growth}}$ | $N_{\text{fam once selfed}}$ |
| --- | --- | --- | --- | --- | --- | --- | --- | --- |
| 1 | TN | Corn | 35.775237 | -85.903419 | 8 | 31 | 10 | 4 |
| 2 | NC | Corn | 34.595714 | -77.927484 | 79 | 387 | 60 |  |
| 4 | NC | Corn | 34.556672 | -79.125602 | 39 | 178 | 79 |  |
| 5 | SC | Soy | 33.859875 | -79.909072 | 34 | 146 | 27 |  |
| 8 | SC | Corn | 34.297195 | -79.991259 | 40 | 180 | 73 | 8 |
| 9 | NC | Soy | 34.924044 | -77.796171 | 40 | 189 | 46 | 8 |
| 10 | NC | Soy | 34.983161 | -78.039309 | 28 | 123 | 24 | 8 |
| 11 | NC | Corn | 34.527135 | -78.756704 | 38 | 178 | 81 | 8 |
| 12 | SC | Cotton | 34.145812 | -79.865313 | 42 | 202 | 79 | 8 |
| 14 | NC | Soy | 35.424763 | -77.917121 | 40 | 192 | 76 | 8 |
| 15 | SC | Soy | 34.104209 | -79.073735 | 41 | 193 | 81 | 8 |
| 16 | SC | Alfalfa | 34.10535 | -79.183234 | 41 | 203 | 78 |  |
| 17 | SC | Soy | 34.159155 | -79.272908 | 27 | 135 | 82 |  |
| 18 | SC | Corn | 34.156593 | -79.27027 | 39 | 193 | 79 |  |
| 19 | NC | Corn | 34.508193 | -78.70899 | 35 | 138 | 70 | 8 |
| 20 | TN | Corn | 35.830692 | -85.777871 | 38 | 169 | 13 | 2 |
| 21 | TN | Soy | 35.369816 | -77.877314 | 40 | 200 | 70 |  |
| 22 | NC | Corn | 36.1436 | -78.053422 | 17 | 78 | 69 |  |
| 23 | TN | Corn | 35.067905 | -86.62955 | 40 | 195 | 77 |  |
| 26 | TN | Soy | 35.533413 | -85.951902 | 50 | 250 | 86 |  |
| 28 | SC | Corn | 34.097917 | -80.377715 | 35 | 172 | 80 |  |
| 29 | NC | Corn | 34.705135 | -78.738897 | 30 | 122 | 75 | 7 |
| 30 | TN | Corn | 35.31105 | -85.945003 | 42 | 206 | 82 |  |
| 31 | TN | Corn | 35.608482 | -85.846379 | 31 | 141 | 69 | 8 |
| 32 | TN | Corn | 35.099356 | -86.225509 | 15 | 56 | 33 | 5 |
| 33 | OH | Corn | 39.858763 | -83.669821 | 52 | 260 | 79 |  |
| 34 | OH | Corn | 39.44316 | -83.910189 | 48 | 238 | 82 |  |
| 35 | IN | Corn | 39.853945 | -85.770156 | 28 | 120 | 80 |  |
| 36 | IN | Corn | 40.565608 | -85.503826 | 33 | 146 | 85 |  |
| 37 | OH | Corn | 39.583755 | -83.758264 | 49 | 243 | 79 |  |
| 38 | OH | Soy | 39.515447 | -83.407431 | 46 | 230 | 88 |  |
| 39 | IN | Corn | 39.988984 | -85.742262 | 38 | 188 | 75 |  |
| 40 | OH | Soy | 41.284684 | -83.847252 | 50 | 237 | 84 |  |
| 41 | VA | Corn | 38.636343 | -78.472921 | 41 | 200 | 78 |  |
| 42 | VA | Corn | 38.373523 | -78.662516 | 49 | 230 | 82 | 8 |
| 43 | VA | Soy | 36.886448 | -78.553156 | 24 | 89 | 28 | 2 |
| 44 | VA | Soy | 38.285415 | -78.797088 | 20 | 98 | 75 |  |
| 45 | VA | Soy | 36.847945 | -78.595042 | 42 | 209 | 76 |  |
| 46 | TN | Soy | 35.536019 | -86.17985 | 34 | 100 | 17 | 2 |
| 47 | SC | Soy | 34.282132 | -79.746597 | 41 | 201 | 83 |  |
| 48 | TN | Corn | 35.31653 | -87.35373 | 31 | 151 | 74 | 8 |
| 51 | TN | Soy | 35.533413 | -85.951902 | 24 | 118 | 56 | 8 |
| 52 | OH | Corn | 41.284684 | -83.847252 | 62 | 310 | 88 |  |

**Table S2.** Regression results between resistance and seed quality traits, accounting for geography (latitude and longitude) for field-collected seeds.

| Trait | Population<br>number | Survival at 1.7 kg a.i.<br>/ha |  | Latitude (scaled) |  | Longitude (scaled) |  |
| --- | --- | --- | --- | --- | --- | --- | --- |
| | | *b* | $\chi^2_1$ | *b* | $\chi^2_1$ | *b* | $\chi^2_1$ |
| Seed weight (g) | 43 | -0.0047 | 4.63* | 0.0016 | 7.33** | -0.0010 | 3.91* |
| Final germination | 43 | -4.89 | 28.62*** | 0.6298 | 8.80** | -0.3784 | 4.13* |
| Abnormally<br>germinating seeds | 42 | 4.24 | 32.52*** | -0.1774 | 1.24 | 0.1826 | 1.58 |
| Seeds needing<br>scarification | 43 | 0.83 | 1.77 | -0.4636 | 7.19** | -0.3488 | 5.22* |
| Scarified seeds that<br>germinated | 40 | -5.87 | 8.85** | 0.0061 | 0.0002 | -0.3342 | 0.65 |
| Height (cm) | 43 | -7.39 | 7.26** | -0.3336 | 0.38 | 1.2210 | 4.76* |
| Leaf number | 43 | -1.08 | 17.57*** | -0.0070 | 0.02 | 0.1589 | 9.17** |
| Largest leaf size (Box<br>cox transformed cm) | 43 | -4.40 | 14.04*** | 0.4080 | 2.96 | 0.4642 | 3.81 |

**Table S3.** PCA loadings and variance explained.

| Trait | Factor1 | Factor2 | Factor3 |
| --- | --- | --- | --- |
| Seed weight | 0.09506 | 0.8414 | -0.16627 |
| Percent germination | 0.29276 | 0.82272 | 0.37541 |
| Percent viable dormant seeds | 0.0339 | 0.12801 | 0.92794 |
| Percent abnormally<br>germinating seeds | -0.15651 | -0.81953 | -0.42013 |
| Height | 0.94253 | 0.04143 | 0.05526 |
| Leaf number | 0.86728 | 0.13535 | 0.06511 |
| Largest leaf size | 0.74853 | 0.28353 | 0.00642 |
| Prop. Variance explained | 0.3316 | 0.3105 | 0.1734 |

**Fig S1.**
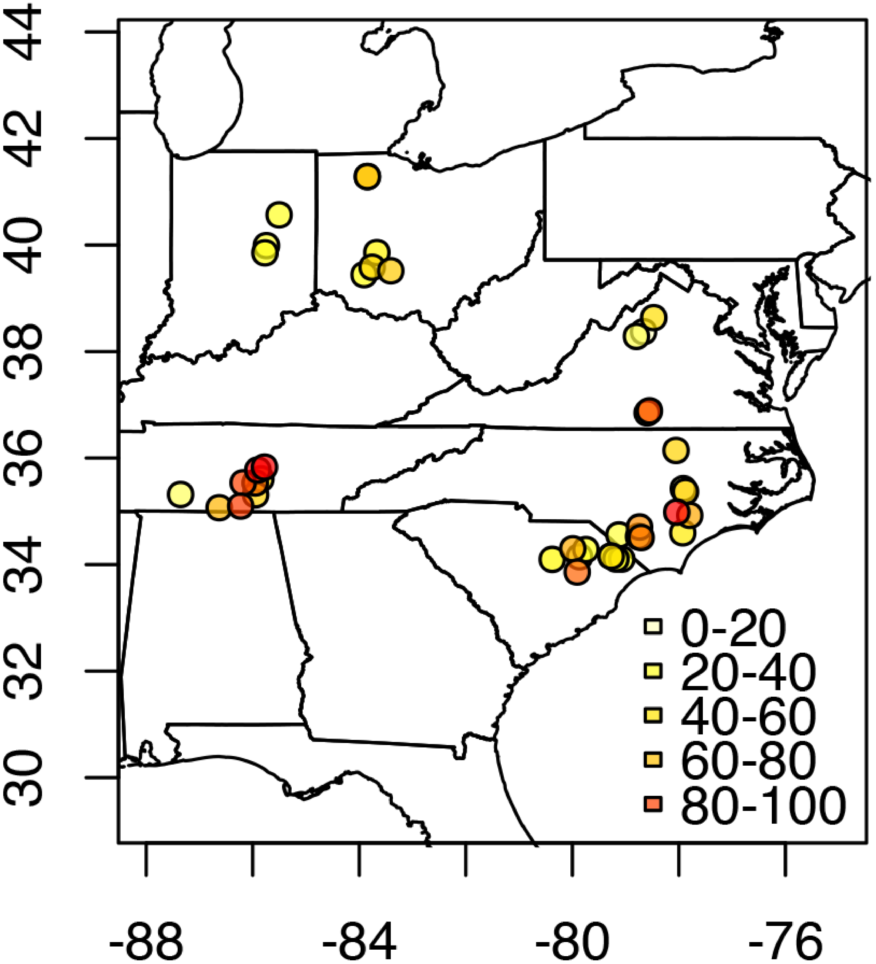
Map of the Eastern US showing the population locations and their survival at 1.7 kg a.i./ha (color of circle)

## References

Abramoff, M. D., P. J. Magelhaes, and S. J. Ram. 2004. Image Processing with ImageJ. Biophotonics International 11:36–42.

Ahrens, W. H. and E. Stoller. 1983. Competition, growth rate, and CO_2_ fixation in triazine-susceptible and-resistant smooth pigweed (*Amaranthus hybridus*). Weed Science 31:438–444.

Alcocer-Ruthling, M., D. C. Thill, and B. Shafii. 1992. Differential competitiveness of sulfonylurea resistant and susceptible prickly lettuce (*Lactuca serriola*). Weed Technology 6:303–309.

Bakker, E. G., C. Toomajian, M. Kreitman, and J. Bergelson. 2006. A genome-wide survey of *R* gene polymorphisms in *Arabidopsis*. The Plant Cell 18:1803–1818.

Barrett, S. H. 1983. Crop mimicry in weeds. Economic Botany 37:255–282.

Baucom, R. S. and R. Mauricio. 2004. Fitness costs and benefits of novel herbicide tolerance in a noxious weed. Proceedings of the National Academy of Sciences of the United States of America 101:13386–13390.

Baucom, R. S. and R. Mauricio. 2008. Constraints on the evolution of tolerance to herbicide in the common morning glory: resistance and tolerance are mutually exclusive. Evolution 62:2842–2854.

Bergelson, J. and C. B. Purrington. 1996. Surveying patterns in the cost of resistance in plants. American Naturalist 148:536–558.

Brabham, C. B., C. K. Gerber, and W. G. Johnson. 2011. Fate of glyphosate-resistant giant ragweed (*Ambrosia trifida*) in the presence and absence of glyphosate. Weed Science 59:506–511.

Brechu-Franco, A., R. Ponce-Salazar, G. Laguna-Hernandez, and J. Marquez-Guzman. 2000. Effect of thermal storage on seed coat dormancy and germination of *Ipomoea purpurea* (L.) Roth (Convolvulaceae) seeds. Phyton 67:187–194.

Christopher, J. T., S. B. Powles, and J. A. Holtum. 1992. Resistance to acetolactate synthase-inhibiting herbicides in annual ryegrass (*Lolium rigidum*) involves at least two mechanisms. Plant Physiology 100:1909–1913.

Christopher, J. T., S. B. Powles, D. R. Liljegren, and J. A. Holtum. 1991. Crossresistance to herbicides in annual ryegrass (*Lolium rigidum*) II. chlorsulfuron resistance involves a wheat-like detoxification system. Plant Physiology 95:1036–1043.

Christopher, J. T., C. Preston, and S. B. Powles. 1994. Malathion antagonizes metabolism-based chlorsulfuron resistance in *Lolium rigidum*. Pesticide Biochemistry and Physiology 49:172–182.

Cousens, R., G. Gill, and E. J. Speijers. 1997. Comment: number of sample populations required to determine the effects of herbicide resistance on plant growth and fitness. Weed Research 37:1–4.

Coustau, C. and C. Chevillon. 2000. Resistance to xenobiotics and parasites: can we count the cost? Trends in Ecology & Evolution 15:378–383.

Culpepper, A. S. 2006. Glyphosate-induced weed shifts. Weed Technology 20:277–281.

Darmency, H., Y. Menchari, V. Le Corre, and C. Délye. 2015. Fitness cost due to herbicide resistance may trigger genetic background evolution. Evolution 69:271–278.

Debban, C., S. Matthews, L. Elliot, K. Pieper, A. Wilson, and R. S. Baucom. 2015. An examination of fitness costs of glyphosate resistance in the common morning glory, *Ipomoea purpurea*. Ecology and Evolution 5:5284–5294.

Délye, C., M. Jasieniuk, and V. Le Corre. 2013a. Deciphering the evolution of herbicide resistance in weeds. Trends in Genetics 29:649–658.

Délye, C., Y. Menchari, S. Michel, É. Cadet, and V. Le Corre. 2013b. A new insight into arable weed adaptive evolution: mutations endowing herbicide resistance also affect germination dynamics and seedling emergence. Annals of botany 111:681–691.

Délye, C., S. Michel, A. Bérard, B. Chauvel, D. Brunel, J. P. Guillemin, F. Dessaint, and V. Le Corre. 2010. Geographical variation in resistance to acetyl - coenzyme A carboxylase - inhibiting herbicides across the range of the arable weed *Alopecurus myosuroides* (black - grass). New Phytologist 186:1005–1017.

Dinelli, G., I. Marotti, P. Catizone, S. Bosi, A. Tanveer, R. Abbas, and D. Pavlovic. 2013. Germination ecology of *Ambrosia artemisiifolia* L. and *Ambrosia trifida* L. biotypes suspected of glyphosate resistance. Open Life Sciences 8:286–296.

El-Kassaby, Y. A., I. Moss, D. Kolotelo, and M. Stoehr. 2008. Seed germination: mathematical representation and parameters extraction. Forest Science 54:220–227.

Fenner, M. 1991. The effects of the parent environment on seed germinability. Seed Science Research 1:75–84.

Fernandez-Cornejo, J., R. Nehring, C. Osteen, S. Wechsler, A. Martin, and A. Vialou. 2014. Pesticide use in U.S. agriculture: 21 selected crops, 1960-2008, EIB-124. U.S. Department of Agriculture, Economic Research Service.

Forcella, F., R. L. B. Arnold, R. Sanchez, and C. M. Ghersa. 2000. Modeling seedling emergence. Field Crops Research 67:123–139.

Galloway, L. F. 2001. The effect of maternal and paternal environments on seed characters in the herbaceous plant *Campanula americana* (Campanulaceae). American Journal of Botany 88:832–840.

Gemmill, A. W. and A. F. Read. 1998. Counting the cost of disease resistance. Trends in Ecology & Evolution 13:8–9.

Giacomini, D., P. Westra, and S. M. Ward. 2014. Impact of genetic background in fitness cost studies: an example from glyphosate-resistant *Palmer amaranth*. Weed Science 62:29–37.

Glettner, C. E. and D. E. Stoltenberg. 2015. Noncompetitive growth and fecundity of Wisconsin giant ragweed resistant to glyphosate. Weed Science 63:273–281.

Goh, S. S., M. M. Vila-Aiub, R. Busi, and S. B. Powles. 2015. Glyphosate resistance in *Echinochloa colona:* phenotypic characterisation and quantification of selection intensity. Pest management science.

Gould, F. 1995. Comparisons between resistance management strategies for insects and weeds. Weed Technology:830–839.

Hartl, D. L. and A. G. Clark. 1989. Principles of population genetics. Sinauer associates, Sunderland, Mass.

Heap, I. 2015. The International survey of herbicide resistant weeds.

Holt, J. 1988. Reduced growth, competitiveness, and photosynthetic efficiency of triazine-resistant *Senecio vulgaris* from California. Journal of Applied Ecology 25:307–318.

Ismail, B., T. Chuah, S. Salmijah, Y. Teng, and R. Schumacher. 2002. Germination and seedling emergence of glyphosate - resistant and susceptible biotypes of goosegrass (*Eleusine indica* [L.] Gaertn.). Weed Biology and Management 2:177–185.

Kuester, A., S. M. Chang, and R. S. Baucom. 2015. The geographic mosaic of herbicide resistance evolution in the common morning glory, *Ipomoea purpurea:* Evidence for resistance hotspots and low genetic differentiation across the landscape. Evolutionary Applications 8:821–833.

Leslie, T. and R. S. Baucom. 2014. De novo assembly and annotation of the transcriptome of the agricultural weed *Ipomoea purpurea* uncovers gene expression changes associated with herbicide resistance. G3: Genes | Genomes | Genetics 4:2035–2047.

Lewontin, R. C. 1974. The genetic basis of evolutionary change. Columbia University Press, New York.

Menchari, Y., C. Camilleri, S. Michel, D. Brunel, F. Dessaint, V. Le Corre, and C. Délye. 2006. Weed response to herbicides: regional - scale distribution of herbicide resistance alleles in the grass weed *Alopecurus myosuroides*. New Phytologist 171:861–874.

Menchari, Y., B. Chauvel, H. Darmency, and C. Délye. 2008. Fitness costs associated with three mutant acetyl - coenzyme A carboxylase alleles endowing herbicide resistance in black - grass *Alopecurus myosuroides*. Journal of Applied Ecology 45:939–947.

Mortimer, A. M. 1997. Phenological adaptation in weeds—an evolutionary response to the use of herbicides? Pesticide Science 51:299–304.

Neve, P., R. Busi, M. Renton, and M. M. Vila-Aiub. 2014. Expanding the eco - evolutionary context of herbicide resistance research. Pest Management Science 70:1385–1393.

O'Donovan, J., J. Newman, R. Blackshaw, K. Harker, D. Derksen, and A. Thomas. 1999. Growth, competitiveness, and seed germination of triallate/difenzoquatsusceptible and-resistant wild oat populations. Canadian Journal of Plant Science 79:303–312.

Owen, M. J., D. E. Goggin, and S. B. Powles. 2014. Intensive cropping systems select for greater seed dormancy and increased herbicide resistance levels in *Lolium rigidum* (annual ryegrass). Pest Management Science.

Paris, M., F. Roux, A. Berard, and X. Reboud. 2008. The effects of the genetic background on herbicide resistance fitness cost and its associated dominance in *Arabidopsis thaliana*. Heredity 101:499–506.

Pedersen, B. P., P. Neve, C. Andreasen, and S. B. Powles. 2007. Ecological fitness of a glyphosate-resistant *Lolium rigidum* population: Growth and seed production along a competition gradient. Basic and Applied Ecology 8:258–268.

Pilling, J., L. Willmitzer, and J. Fisahn. 2000. Expression of a *Petunia inflata* pectin methyl esterase in *Solanum tuberosum* L. enhances stem elongation and modifies cation distribution. Planta 210:391–399.

Platenkamp, G. A. and R. G. Shaw. 1993. Environmental and genetic maternal effects on seed characters in *Nemophila menziesii*. Evolution 47:540–555.

Powles, S. B. and Q. Yu. 2010. Evolution in action: plants resistant to herbicides. Annual Review of Plant Biology 61:317–347.

Preston, C. and S. B. Powles. 1998. Amitrole inhibits diclofop metabolism and synergises diclofop-methyl in a diclofop-methyl-resistant diotype of *Lolium rigidum*. Pesticide Biochemistry and Physiology 62:179–189.

Primack, R. B. and H. Kang. 1989. Measuring fitness and natural selection in wild plant populations. Annual Review of Ecology and Systematics 20:367–396.

Rausher, M. D. and E. L. Simms. 1989. The evolution of resistance to herbivory in *Ipomoea purpurea*. I. Attempts to detect selection. Evolution 43:563–572.

Ren, C. and A. R. Kermode. 2000. An increase in pectin methyl esterase activity accompanies dormancy breakage and germination of yellow cedar seeds. Plant Physiology 124:231–242.

Roach, D. A. and R. D. Wulff. 1987. Maternal effects in plants. Annual Review of Ecology and Systematics 18:209–235.

Schmitt, J., J. Niles, and R. D. Wulff. 1992. Norms of reaction of seed traits to maternal environments in *Plantago lanceolata*. American Naturalist 139:451–466.

Shrestha, A., K. M. Steinhauer, M. L. Moretti, B. D. Hanson, M. Jasieniuk, K. J. Hembree, and S. D. Wright. 2014. Distribution of glyphosate-resistant and glyphosate-susceptible hairy fleabane (*Conyza bonariensis*) in central California and their phenological development. Journal of Pest Science 87:201–209.

Simms, E. L. and M. D. Rausher. 1987. Costs and benefits of plant resistance to herbivory. American Naturalist 130:570–581.

Simms, E. L. and M. D. Rausher. 1989. The evolution of resistance to herbivory in *Ipomoea purpurea*. II. Natural selection by insects and costs of resistance. Evolution 43:573–585.

Stahl, E. A., G. Dwyer, R. Mauricio, M. Kreitman, and J. Bergelson. 1999. Dynamics of disease resistance polymorphism at the *Rpm1* locus of *Arabidopsis*. Nature 400:667–671.

Strauss, S. Y., J. A. Rudgers, J. A. Lau, and R. E. Irwin. 2002. Direct and ecological costs of resistance to herbivory. Trends in Ecology & Evolution 17:278–285.

Tardif, F. J., I. Rajcan, and M. Costea. 2006. A mutation in the herbicide target site acetohydroxyacid synthase produces morphological and structural alterations and reduces fitness in *Amaranthus powellii*. New Phytologist 169:251–264.

Vila-Aiub, M. M., S. S. Goh, T. A. Gaines, H. Han, R. Busi, Q. Yu, and S. B. Powles. 2014. No fitness cost of glyphosate resistance endowed by massive EPSPS gene amplification in *Amaranthus palmeri*. Planta 239:793–801.

Vila-Aiub, M. M., P. Neve, and S. B. Powles. 2009a. Evidence for an ecological cost of enhanced herbicide metabolism in *Lolium rigidum*. Journal of Ecology 97:772–780.

Vila-Aiub, M. M., P. Neve, and S. B. Powles. 2009b. Fitness costs associated with evolved herbicide resistance alleles in plants. New Phytologist 184:751–767.

Vila-Aiub, M. M., P. Neve, and F. Roux. 2011. A unified approach to the estimation and interpretation of resistance costs in plants. Heredity 107:386–394.

Vila-Aiub, M. M., P. Neve, K. Steadman, and S. Powles. 2005. Ecological fitness of a multiple herbicide - resistant *Lolium rigidum* population: dynamics of seed germination and seedling emergence of resistant and susceptible phenotypes. Journal of Applied Ecology 42:288–298.

Vogwill, T., M. Lagator, N. Colegrave, and P. Neve. 2012. The experimental evolution of herbicide resistance in *Chlamydomonas reinhardtii* results in a positive correlation between fitness in the presence and absence of herbicides. Journal of Evolutionary Biology 25:1955–1964.

Weaver, S. and S. Warwick. 1982. Competitive relationships between atrazine resistant and susceptible populations of *Amaranthus retroflexus* and *A. powellii* from southern Ontario. New Phytologist 92:131–139.

Weaver, S. E. and P. B. Cavers. 1979. The effects of date of emergence and emergence order on seedling survival rates in *Rumex crispus* and *R obtusifolius*. Canadian Journal of Botany 57:730–738.

Webster, T. M. and G. E. MacDonald. 2001. A survey of weeds in various crops in Georgia 1. Weed Technology 15:771–790.

Williams, M. M., N. Jordan, and C. Yerkes. 1995. The fitness cost of triazine resistance in jimsonweed (*Datura stramonium* L.). American Midland Naturalist:131–137.

Yu, Q., A. Collavo, M.-Q. Zheng, M. Owen, M. Sattin, and S. B. Powles. 2007. Diversity of acetyl-coenzyme A carboxylase mutations in resistant *Lolium* populations: evaluation using clethodim. Plant Physiology 145:547–558.

Yu, Q., H. Han, and S. B. Powles. 2008. Mutations of the ALS gene endowing resistance to ALS - inhibiting herbicides in *Lolium rigidum* populations. Pest Management Science 64:1229–1236.

Yu, Q., H. Han, M. M. Vila-Aiub, and S. B. Powles. 2010. AHAS herbicide resistance endowing mutations: effect on AHAS functionality and plant growth. Journal of Experimental Botany 61:3925–3934.

